# CRISPR/Cas13-mediated Dynamin 2 reduction therapy in a canine model of *DNM2*-related centronuclear myopathy

**DOI:** 10.1101/2025.09.05.672814

**Authors:** Andréa Carlier, Inès Barthélémy, Nicolas Blanchard-Gutton, Sophie Chateau-Joubert, Frédéric Auradé, Teoman Ozturk, Nathalie Didier, Frédéric Relaix, Laurent Tiret, Stéphane Blot, Isabel Punzón

## Abstract

We determined the potential of CRISPR/Cas13 technology as a therapeutic approach for centronuclear myopathies (CNMs) by reducing the expression of a single protein, DNM2. CNMs are severe congenital rare muscle disorders that result in muscle hypotrophy and weakness, with no cure. CNMs frequently result from mutations in either *BIN1*, *MTM1*, or *DNM2* genes, with DNM2 being a key GTPase that plays a pivotal role in muscle membrane interactions with MTM1 and BIN1. Previous studies indicate that reducing *DNM2* transcript expression by half could correct CNM phenotypes regardless the genetic forms, paving the way for a broad-spectrum CNM-therapy. We evaluated CRISPR/Cas13X.1-mediated *DNM2* transcript knockdown, as a therapeutic application in a unique naturally-occurring canine CNM model harboring the *DNM2^R465W^*^/+^ mutation, the most frequent pathogenic variant in patients. We show that *in vivo* intramuscular AAV-mediated CRISPR/Cas13X.1 injections, led to a reduction in *DNM2* transcript and protein levels at one and two months post-treatment. Our results demonstrate the feasibility of CRISPR/Cas13-based therapy for CNM in a large animal model, paving the way for advancing this approach towards clinical trials.

## INTRODUCTION

Centronuclear myopathies (CNM) are rare congenital neuromuscular diseases and are mainly characterized by hypotrophy and muscle weakness with centralized nuclei in the fibers [1]. Affected patients are mostly newborns and children at disease onset, that present a heterogeneous range of disease severity [2, 3]. Three main forms of CNM have been classified depending on the genes involved and their mode of inheritance. X-linked myotubular myopathy (*XLMTM1*) is the most common CNM disease and is caused by mutations in the *Phosphoinositide phosphatase myotubularin* gene (*MTM1*) and corresponds to a severe form with an average short lifespan [4]. Autosomal dominant mutations in the *Dynamin2* gene (*DNM2*) give rise to moderate forms of CNM [5, 6]. The severity of *DNM2*-CNM is variable with different onset ages [7]. Autosomal recessive mutations in the membrane remodeling protein *Amphiphysin2* gene (*BIN1*) can be severe with early onset and some individuals may have milder forms of the disease [1, 8]. Mutations in other genes have also been indicated in CNM myopathies, including genes encoding for Titin and Ryanodine receptor [9, 10]. All these proteins interact within the muscle as a network and are implicated in muscle fiber membrane stability [11]. Different therapeutic approaches are considered for CNMs from pharmacological to cell and gene therapies, but there is no cure to date [12–14].

The DNM2 protein is a large ubiquitous GTPase implicated in membrane remodeling processes, including endocytosis, membrane traffic and regulation of microtubules and the actin cytoskeleton [15, 16]. This GTPase plays an important role in membrane fission and vesicle formation. The R465W mutation is the most commonly found in patients suffering from *DNM2* related forms of CNM with slowly progressing myopathy, muscle weakness and ptosis [17]. This mutation lies in the middle domain of the protein, flanking the GTPase domain and has been described as a gain-of-function mutation of *DNM2* [18]. Knock-in mice harboring the R465W mutation developed progressive muscle atrophy, decreased muscle strength and structural disorganization in muscle fibers [19]. It was recently demonstrated that DNM2 was upregulated in the *DNM2*-unrelated CNMs, and that its overexpression induced a CNM phenotype in mice [20]. The reduction of *DNM2* rescues the *DNM2*-related CNM [13]. The same group described that a reduction of 50% of DNM2 (by crossing with heterozygous *Dnm2* mice) led to a rescued phenotype of *Xlmtm1* mouse muscles [21]. *DNM2* downregulation also rescued the *BIN1* phenotype in mice [22]. The use of antisense oligonucleotides (ASO) and shRNA in mice to diminish DNM2 protein has proven effective opening the door to a pan-therapy for several CNM diseases by the regulation of DNM2 protein expression [23]. Most proposed therapies are focused on allele inactivation using antisense oligonucleotides and shRNAs. A phase 1/2 clinical trial to evaluate the efficacy of a strategy of degradation of *DNM2* mRNAs via the antisense oligonucleotide technology was stopped in spite of very encouraging therapeutic response, due to adverse events on treated patients (ClinicalTrials.gov Identifier: NCT04033159). The clinical trial for *MTM1* gene transfer was placed on hold due to serious adverse effects as well (ClinicalTrials.gov Identifier: NCT03199469) [24]. The adverse effects seem to be due to liver cholestasis related to MTM1 absence. Recently it has been also shown in a zebrafish model of *XLMTM1* that reduction of DNM2 improved the hepatic phenotype, presumably due to defects in the bile flux [25]. DNM2 regulation appears to be key target for CNMs phenotype improvement. It is therefore necessary to continue the investigation of alternative strategies, effective and without adverse effects, in the hope that one of them can be rapidly proposed to patients.

It is worth noting that spontaneous cases of CNM have occurred in dogs, which closely resemble the human disease, facilitating efficient translational research as it was the case with *MTM1* Labrador Retriever dog model that allowed the validation of a gene therapy clinical trial [24, 26, 27]. A *BIN1* mutation was also found in a Great Dane dog and a *HACD1* in a Labrador Retriever [28, 29]. Our group has recently obtained and characterized a new canine model of CNM disease, the *DNM2^R465W/+^* dog [30]. The colony was developed from a male Border Collie that spontaneously presented the most common human (30% of patients) *DNM2* mutation: the R465W [18]. Comprehensive characterization of these animals was performed and altogether allowed us to confirm the mutation is fully penetrant and leads to a slowly progressive CNM in dogs resembling the human disease in several aspects. From the age of two months the first signs of disease were seen at the histological level, with oxidative staining revealing necklace fibers with central clustering of mitochondria [31].

In the new era of genome editing, the CRISPR/Cas system has emerged as a powerful tool, with clinical trials currently underway for rare diseases [32]. Recently the efficacy of CRISPR/Cas9 for genome editing in *Dnm2^R465W/+^* human and mouse myoblasts has been proven by the rescue of the CNM-phenotype [33]. This study pointed out the potential risk of off-targets associated with this genome editing tool, its very low efficiency and the fact that homologous direct recombination cannot occur in post-mitotic cells, such as muscle fibers, which is a drawback for *in vivo* applications. Recently, a new Cas protein has been identified in the Class 2 CRISPR/Cas system, known as Cas13 that binds RNA instead of DNA substrates [34, 35]. Several studies showed the possibilities of the discovered Cas13 protein for RNA editing purposes. This protein, which is found in bacteria, functions as an RNA-guided RNAse, providing protection against RNA-phage infection. It achieves this by cleaving single-strand (ss) RNA molecules through a short CRISPR guide RNA (gRNA) that is complementary to the target sequence. Cas13 stands out for its specificity for RNA, and opens up new perspectives in molecular biology, as RNA editing makes it possible to modulate gene expression without altering the genome [36, 37]. Further developments have led to the discovery of more efficient and compact versions of Cas13, broadening the range of possible applications (RNA knockdown, RNA repair system, or RNA imaging) [36, 38, 39]. Notably, the CRISPR/Cas13 RNA-editing therapy has recently entered its first clinical trial (ClinicalTrials.gov Identifier: NCT06031727), making a significant step toward therapeutic use.

Given the urgent need for CNM therapies and the promise of DNM2 as a unique therapeutic target modulation, the objective of this study was to ascertain the therapeutic potential of CRISPR/Cas13-mediated *DNM2* transcript knockdown. We demonstrated that a targeted ∼50% reduction in *DNM2* mRNA levels could be achieved *in vitro*, leading to a consistent decrease in DNM2 protein and an inversion of the pathological phenotype. Building on these promising results, we further validated the strategy *in vivo*. Local intra-muscular AAV-mediated delivery of the CRISPR/Cas13X.1 system into our clinically relevant canine CNM model, successfully reduced *DNM2* mRNA and protein levels within the treated muscle bundles, underscoring its therapeutic potential and the relevance to further develop this approach.

## RESULTS

### Oxidative and autophagy marker dysregulation in the *DNM2^R465W^*^/+^ dog muscles

Our team has previously characterized the histological features of muscle biopsies from the *DNM2^R465W/+^* dog, demonstrating hallmark signs of centronuclear myopathy (CNM). These observations mainly revealed prominent structural alterations within the cytoplasm, including abnormal accumulations of oxidative material (detectable using H&E, oxidative, and COX stains). These accumulations were predominantly located in the central region of myofibres [31]. Given the established function of the DNM2 protein in the autophagy process [40, 41], we hypothesized that the *DNM2^R465W^*^/+^ mutation would impact muscle fiber organization. A study was conducted on the expression levels of other proteins, in addition to DNM2 itself. These included LC3 (microtubule-associated protein light chain 3), p62 (a selective autophagy receptor) as autophagy marker proteins, and COX (Cytochrome C). Immunofluorescence (IF) of p62 (Fig.1A) and staining for COX [31] revealed a reorganization and accumulation detected in the center of the myofibers. Western blot (WB) analysis confirmed a significant upregulation of DNM2 and p62 protein levels in *DNM2^R465W^*^/+^ muscles compared to controls, along with a clear trend towards increased COX expression (Fig.1B-E).

**Fig.1:**
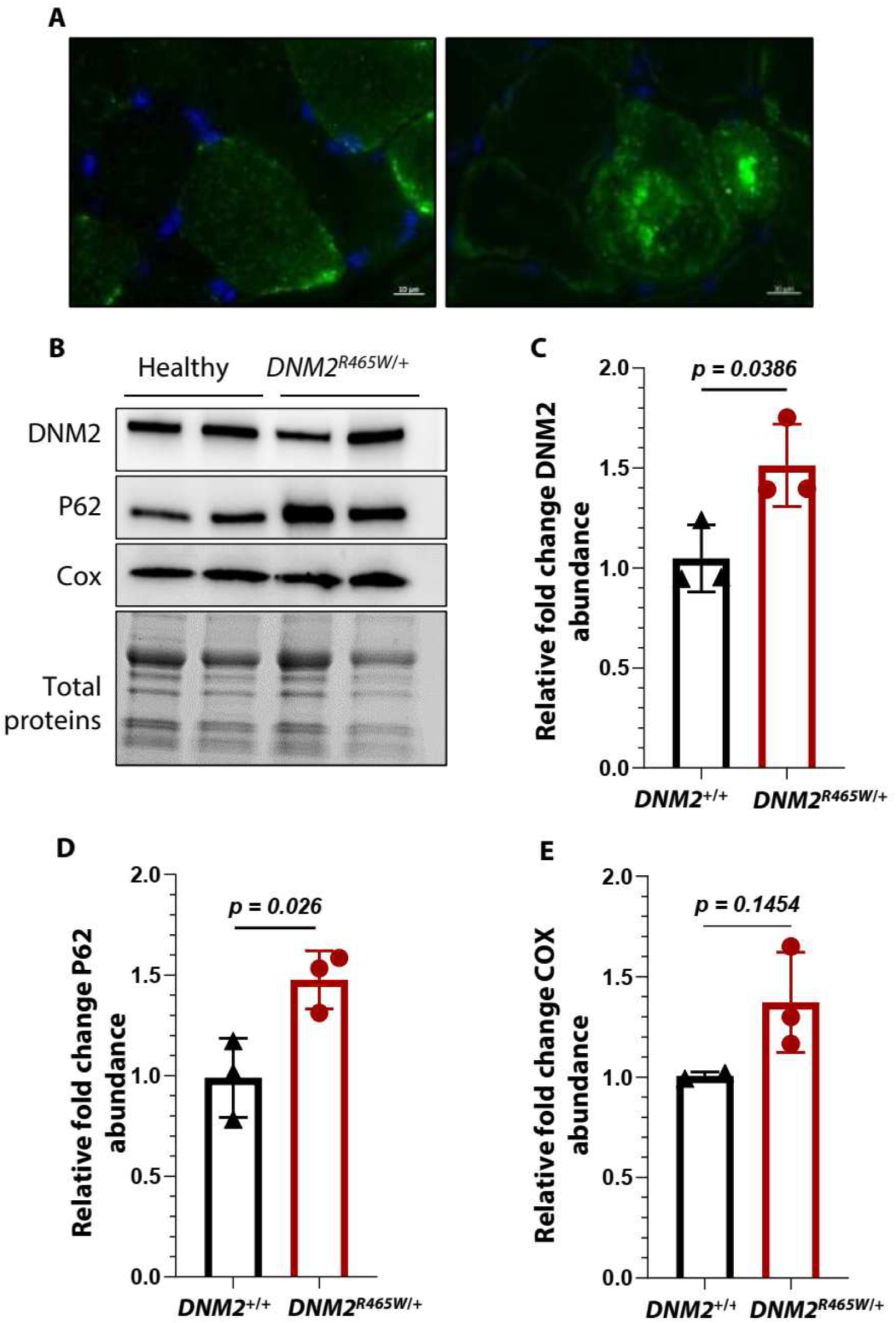
Defects at the histological level of canine *DNM2^R465W/+^* model muscles. (A) Immunofluorescence (IF) staining for p62 on a healthy (left) or *DNM2^R465W^*^/+^ (right) *Biceps femoris* muscles at 12 months of age. Nuclei are counterstained with DAPI. Scale bar: 10µm. (**B**) Representative WB of DNM2, p62 and COX proteins from cryo-biopsies from BF muscle of three *DNM2^R465W/^*^+^ dogs (36 months-old), their control healthy littermate and one/two additional *DNM2*^+/+^ controls. (**B**) Quantification of the intensity of Western blot protein bands intensity of (**C**) DNM2, (**D**) p62 and (**E**) Cytochrome C (COX). Values were normalized to values of *DNM2^+^*^/+^ controls. Data are show as means ± SEM per dogs; Unpaired t-test with Welch correction.

### *DNM2^R465W^*^/+^ cells *in vitro* studies highlight quantifiable alterations in GTPase activity, endocytosis and autophagy flux

We sought to determine whether the histological impairments in myofibers are cell autonomous, and to address this question we obtained *DNM2^R465W^*^/+^ primary myoblasts from muscle biopsies. A number of key characteristics related to the DNM2 protein function were analyzed, with a view to use the most relevant of them to quantify the reversion of the cell phenotype by the CRISPR-Cas13 system. They included assessments of myoblast fusion capacity, GTPase activity, endocytosis, and autophagy activity.

First, we assessed the overall expression and localization of DNM2 protein in diseased cells compared to healthy cells by an IF labelling. The DNM2 protein was expressed in *DNM2^R465W/+^* myoblasts (Fig. 2A). In order to study the potential impact of the mutation on fusion capacity, primary cells were cultured in a differentiation medium for seven days. Our research has revealed that *DNM2^R465W/+^* myoblasts maintain the capacity to form myotubes in a manner consistent with healthy cells (Fig. 2B). The Dynamin 2 protein is a large GTPase and the R465W mutation appears to increase the activity of this protein by stabilizing the oligomerization of the protein for the GTPase function [42, 43]. To evaluate the effect of the mutation on the overall GTPase activity of myoblasts, we used a colorimetric reaction to measure the GTPase activity. *DNM2^R465W^*^/+^ cells showed increased relative GTPase activity in comparison to healthy ones. Experiments were duplicated with the addition of Dynasore [44, 45], an inhibitor of Dynamin GTPase activity. *DNM2^R465W/+^* yielded an elevated GTPase activity compared to *DNM2^+/+^* cells, that could be attributed to Dynamins, as treatment with the inhibitor resulted in similar levels of reduced GTPase activity regardless of the genotype of cells (Fig. 2C). In light of the role of DNM2 in vesicle fission during Clathrin-mediated endocytosis [46, 47], transferrin uptake was measured in myoblasts. The uptake of cy3-labelled transferrin was measured by flow cytometry, and the results showed an increase in transferrin uptake by *DNM2^R465W^*^/+^ cells compared to the controls (Fig. 2D). The Dynasore inhibitor was used in this experiment, showing a difference between healthy and mutated cells, *DNM2^R465W^*^/+^ cells appear to be less sensitive to the inhibitor, resulting in no reduction in transferrin uptake.

**Fig. 2:**
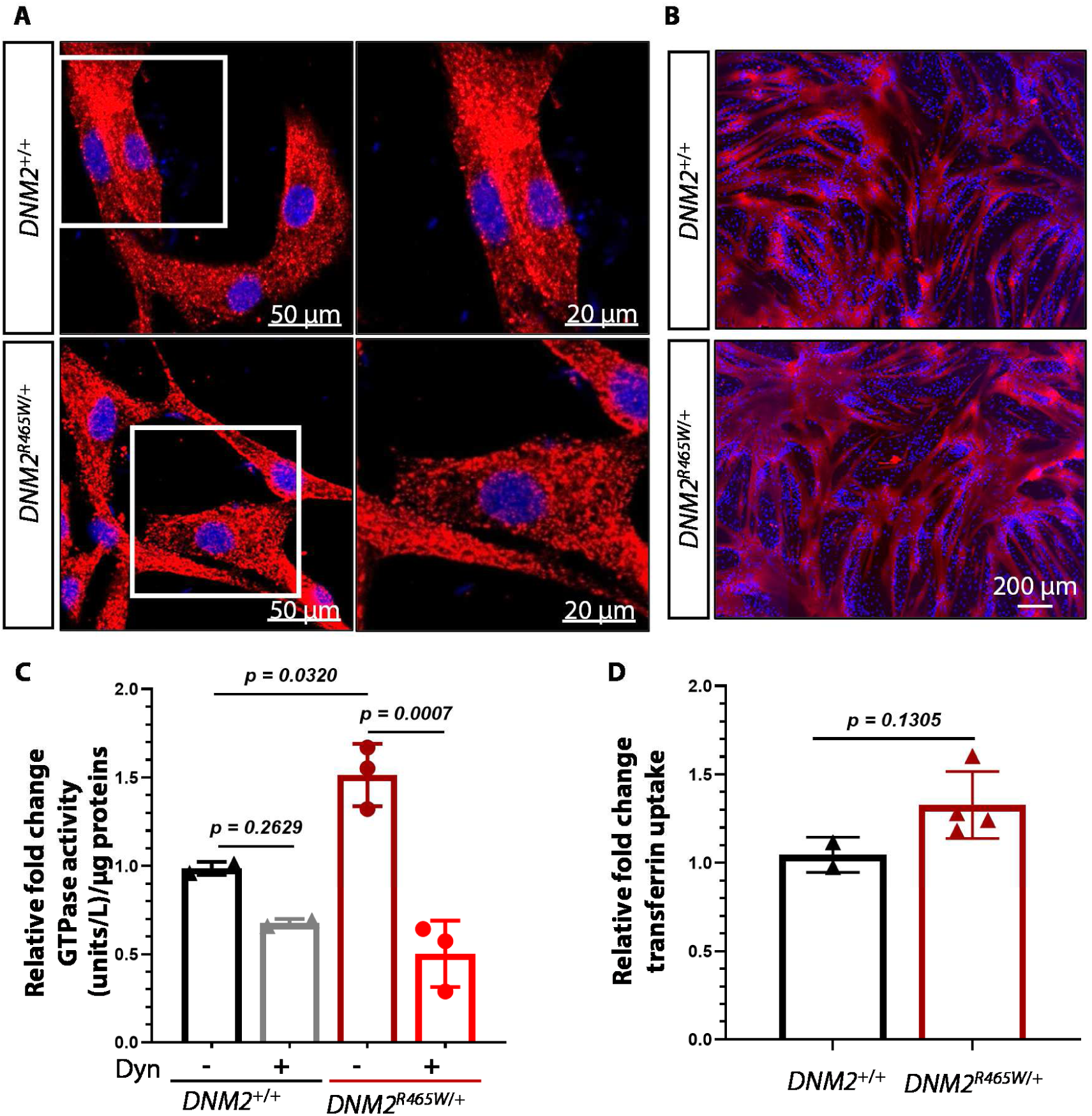
*DNM2^R465W^*^/+^ primary myoblasts maintain fusion capacity but show increased GTPase activity and endocytosis. (**A**) Immunofluorescence (IF) staining for Dynamin 2 (DNM2) on *DNM2^+/+^* (top panel) or *DNM2^R465W^*^/+^ (bottom panel) primary myoblasts. Nuclei are counterstained with DAPI. (**B**) IF staining for Myosin Heavy Chain (MHC) on *DNM2*^+/+^ (top panel) and *DNM2^R465W^*^/+^ (bottom panel) differentiating myoblasts. Nuclei are counterstained with DAPI. (**C**) GTPase activity in primary myoblasts from muscle biopsies of three *DNM2^R465W/^*^+^ dogs, with or without addition of the Dynamin inhibitor, Dynasore (Dyn); normalized to values of two *DNM2*^+/+^ control myoblasts. (**D**) Quantification of Cyanine3-labelled transferrin uptake in primary myoblasts from muscle biopsies of four *DNM2^R465W^*^/+^ dogs; normalized to values of two *DNM2*^+/+^ control myoblasts. Data are shown as means ± SEM per dogs; One-way ANOVA with Tukey’s post hoc test (C) and unpaired t-test with Welch correction.

DNM2 has been demonstrated to play a role in autophagy via the reformation of autophagic lysosomes. Autophagy defects were highlighted by IF labeling of LC3 and p62 autophagy-related proteins in *DNM2^R465W^*^/+^ cells. We used a chloroquine (Cq) treatment to block autophagy flux by preventing the degradation of cellular components in lysosomes [48]. Some spot accumulations and the appearance of ‘vacuoles’ were observed in the mutated cells (Fig. 3A). A WB was performed to quantify these proteins, showing a tendency towards a decreased p62 protein level and a significant increase in the LC3 II protein level in cells with or without chloroquine addition, showing differences between *DNM2^R465W^*^/+^ cells and healthy ones (Fig. 3B-C).

**Fig. 3:**
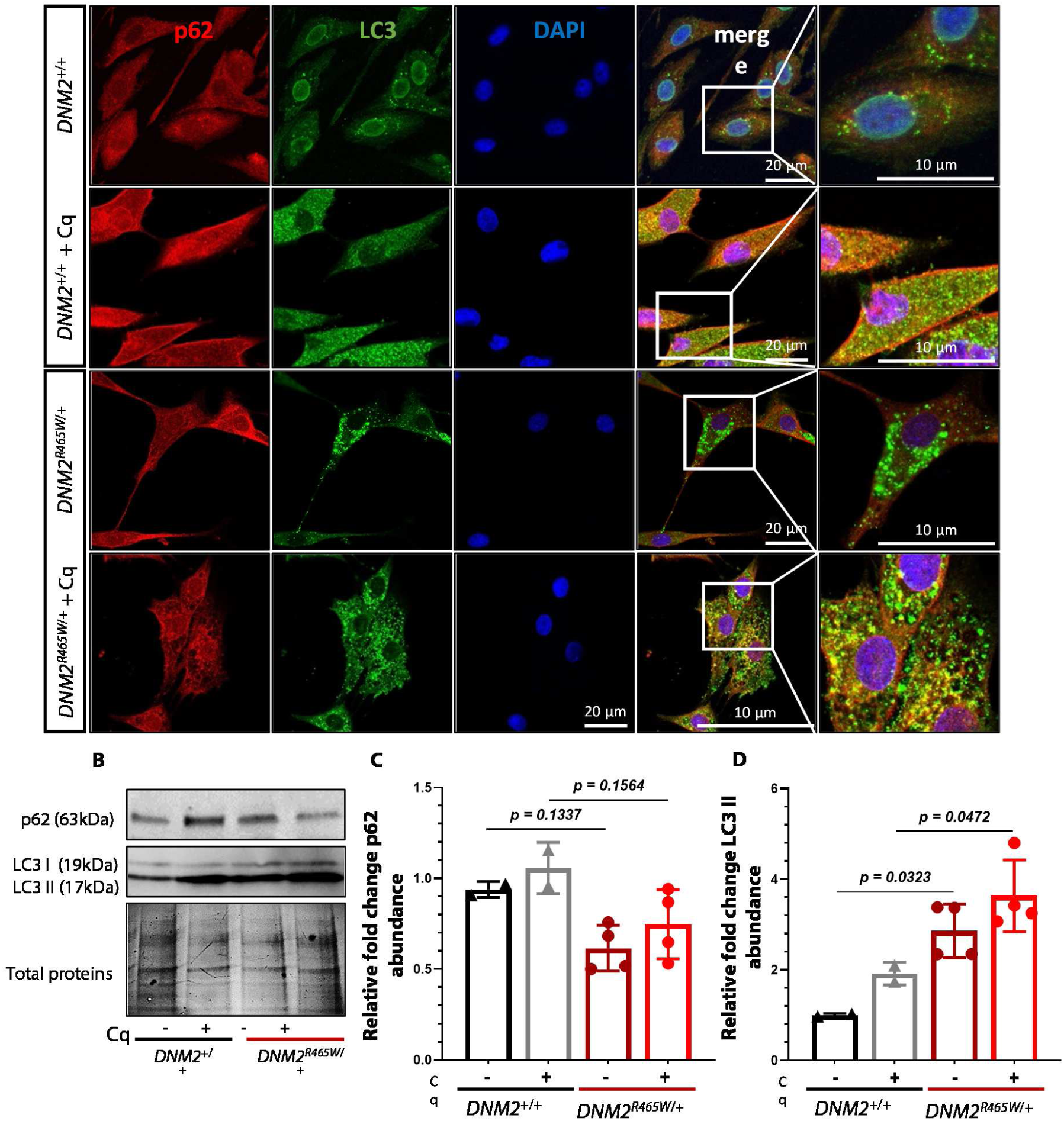
The R465W mutation impairs autophagy in primary myoblasts. (**A**) IF staining for P62 and LC3 proteins on *DNM2*^+/+^ (top panel) and *DNM2^R465W^*^/+^ (bottom panel) primary myoblasts, with or without addition of chloroquine (Cq) which blocks autophagy (50µ M for 16 hours). Nuclei are counterstained with DAPI. (**B**) Representative WB performed on primary cells, with or without addition of Cq, staining for P62 and LC3 I and II. (**C**) Quantification of the intensity of P62 marker protein bands in primary myoblasts from muscle biopsies of four *DNM2^R465W^*^/+^ dogs, with or without Cq; normalized to values of two *DNM2*^+/+^ control dogs. (**D**) Quantification of the intensity of LC3 II marker protein bands in primary myoblasts from muscle biopsies of four *DNM2^R465W^*^/+^ dogs, with or without Cq; normalized to values of two *DNM2*^+/+^ control dogs. Data are shown as means ± SEM per dogs; One-way ANOVA with Tukey’s test.

To overcome the limitation of primary cells renewal, *DNM2^R465W^*^/+^ and *DNM2^+/+^* healthy myoblasts were transduced using a lentiviral vector expressing genes for immortalization, *CDK4* and *TERT*. Transduced cells demonstrated their capacity to proliferate and successfully surpassed the 45^th^ passage. In order to assess the accuracy and consistency of results, all the tests indicated above were first conducted in primary myoblasts and then re-validated in immortalized cells. They maintained the NCAM (CD56) and Desmin myoblast markers, and the *DNM2^R465W^*^/+^ primary cells phenotype related to GTPase activity, endocytosis and autophagy (Fig. S1 and S2). These cells have been used for guide RNA (gRNA) and Cas13 *in vitro* selection.

### *In vitro* knockdown of *DNM2* mRNA by CRISPR/CasX.1

In order to test the efficacy of CRISPR/Cas13 in reducing *DNM2* mRNA levels, several guide gRNAs targeting the *DNM2* transcript were designed and tested *in vitro* with the objective of reducing the mRNA by half (therapeutic downregulation) in cells (Fig. S3A). We conducted a trial to assess the best RNA knockdown efficiency using distinct Cas13 variants (LwaCas13a, RfxCas13d, and Cas13X.1) in conjunction with different gRNAs. Transient transfection was performed using a dual plasmid system to express the CRISPR/Cas13. One plasmid expressing gRNAs under the human U6 promoter, and another plasmid driving Cas13 protein expression under a ubiquitous CMV or EF1α promoter. Fluorescent reporter genes (EGFP or mCherry) were used for sorting transfected cells 48 hours post-transfection by flow cytometry. We first started using the LwaCas13a protein with gRNAs cloned into the pC016-LwaCas13a guide expression backbone plasmid (U6 promoter), allowing the expression of a gRNA with a direct repeat sequence (DR) upstream of the guide sequence. For the use of RfxCas13d and Cas13X.1, gRNAs were cloned into the pxr004-RfxCas13d guide expression backbone plasmid (U6 promoter) which flanks the guide with two DR sequences. The results demonstrated that the Cas13X.1 was the most effective achieving a close 50% knockdown of *DNM2* mRNA in *DNM2^R465W^*^/+^ cells (Fig. S3B). Cas13X.1, which has a shorter cDNA sequence, making it easier to clone into an AAV vector, was chosen for subsequent *in vivo* experiments. We demonstrated a *DNM2* transcript reduction with gRNA-11, as well as with the combination of gRNA-11 and gRNA-7, resulting in an effective and reproducible level reduction of 55% and 43%, respectively (Fig. 4A). On this basis, we proceeded to evaluate the combination of Cas13X.1 and these gRNAs. Our aim was to assess whether such mRNA depletion can mitigate the cellular phenotypes previously observed in *DNM2^R465W^*^/+^ cells.

**Fig. 4:**
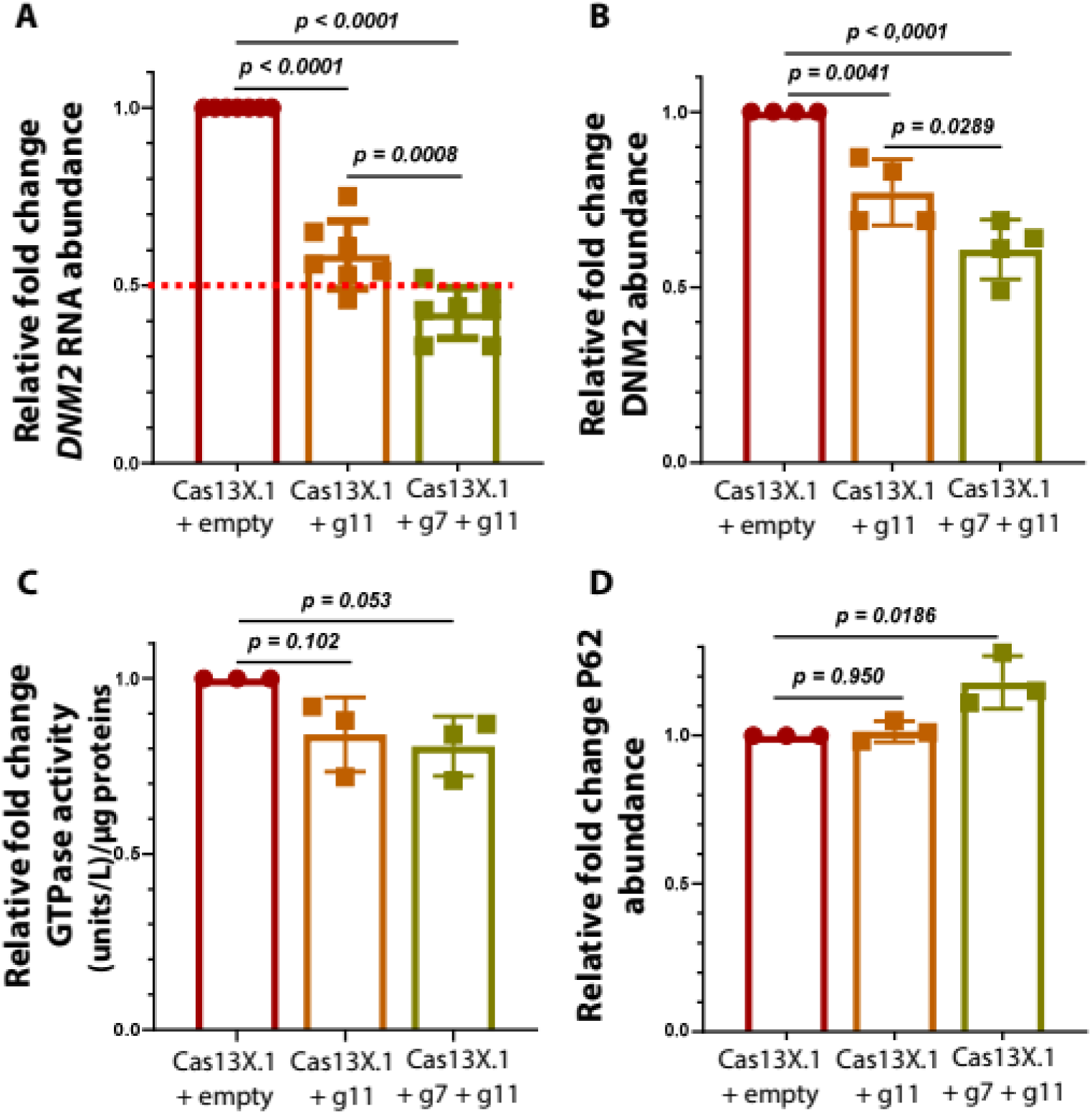
*DNM2^R465W/+^* phenotype reversion through CRISPR/Cas13X.1-mediated *DNM2* mRNA knockdown. (**A**) RT-qPCR performed on RNA extracted from transfected and FACS-sorted cells to quantify *DNM2* mRNA level. Each data point represents an independent biological replicate. *GAPDH* and *RPLS19* used as housekeeping genes and results expressed as the relative fold change abundance of normalized *DNM2* mRNA expression related to the control empty vector (no gRNA) condition. (**B**) Quantification of the intensity of DNM2 protein levels in transfected and FACS-sorted cells; normalized to values of cells transfected with the empty vector condition. (**C**) GTPase activity measured in transfected cells; normalized to values of cells transfected with the empty vector. (**D**) Quantification of the intensity of P62 protein levels in transfected and FACS-sorted cells; normalized to values of cells transfected with the empty vector condition. Each data point on the graph corresponds to an independent experiment. Data are presented as means ± SEM; One-way ANOVA with Tukey’s test.

### CRISPR/Cas13X.1-mediated *DNM2* mRNA reduction reversed phenotype in *DNM2^R465W^*^/+^ cells

We implemented a validation panel composed of conserved and quantifiable readouts: DNM2 protein levels, GTPase enzymatic activity, and p62 protein levels, a marker of autophagy flux.

Western blot analyses of proteins extracted from transfected cells, confirmed that the reduction in *DNM2* transcript levels was paralleled by a proportional decrease in DNM2 protein levels (Fig. 4B). Furthermore, GTPase activity quantification revealed a downward trend in GTPase activity following Cas13X.1-mediated knockdown, consistent with the reduction of the DNM2 protein (Fig. 4C). In addition, p62 levels were also quantified, initially reduced in non-treated cells, showed a slight increase upon DNM2 reduction (Fig. 4D), suggesting a partial restoration of autophagy flux.

### CRISPR/CasX.1 efficiently-mediates *DNM2* mRNA reduction i*n vivo,* improving autophagy-related proteins levels in the three injected dogs

To validate the therapeutic potential of CRISPR/Cas13X.1-mediated *DNM2* knockdown *in vivo,* the Cas13X.1 and the selected gRNAs were packaged in separate AAV-9 vectors (PackGene). An adult *DNM2^R465W^*^/+^ dog was injected intramuscularly with these AAV-9 vectors in the *biceps femoris* (BF) and *triceps brachii* (TB) muscles. Three doses: 3.5×10^10^, 3.5×10^11^ and 3.5×10^12^ vector genomes (vg) were injected in 500 µl of saline in order to select the effective dose. Biopsies were taken at baseline (T0) as a control from each muscle. Then biopsies were collected at one-month (T+1M, BF muscle) and two-month (T+2M, TB muscle) post-injection for molecular and histological analyses of the muscle response to treatment. As early as T+1M, *DNM2* mRNA levels were reduced in treated muscle with the most substantial effect observed at the highest dose at 2 moths post injection (Table 1).

**Table 1.** *In vivo* proof-of-principle: intramuscular injection of AAV-CRISPR/Cas13X.1 shows a reduction in *DNM2* mRNA. Different amounts viral vectors of “AAV-Cas13X.1 + AAV-gRNA7 + AAV-gRNA11”: 3.5 x 10^10^, 3.5 x 10^11^ or 3.5 x 10^12^ viral genome (vg) injected in muscle fascicles. Biopsies were taken at T+0 (before injection), T+1M (1 month) and T+2M (2 months). RT-qPCR analysis of *DNM2* transcripts. The mRNA was referred to *GAPDH* and *RPLS19* housekeeping genes and normalized results are expressed as *DNM2* mRNA abundance related to the biopsies taken at T0.

|  | T + 1M |  |  |  | T + 2M |  |  |  |
| --- | --- | --- | --- | --- | --- | --- | --- | --- |
| vg | T0 | $10^{10}$ | $10^{11}$ | $10^{12}$ | T0 | $10^{10}$ | $10^{11}$ | $10^{12}$ |
| Relative *DNM2* mRNA abundance (fold change) | 1 | 0,92 | 0,735 | 0,775 | 1 | 0,715 | 0,68 | 0,595 |

Two more *DNM2^R465W^*^/+^ canines were similarly injected with the selected dose of 3.5×10¹² vg. Biopsies taken at T0 from the first dog were used as a reference point. To validate the use of T0 biopsies from the first animal as reference controls, we analyzed *DNM2* mRNA levels from both *DNM2^R465W^*^/+^ dogs and their healthy littermate (6–36 months old), revealing minimal inter-individual variability and consistent expression levels (Fig. S4).

The results of the three injected dogs demonstrated a consistent decrease in *DNM2* mRNA from the first month. The reduction was confirmed and appeared to stabilize with reduced variability at two months after injection (Fig. 5A). This decline in mRNA was accompanied by a decrease in DNM2 protein levels at T+2M (Fig. 5B). This outcome demonstrates that the objective of a controlled reduction was therefore met. Moreover, a comparison of the treated muscle with the muscle prior to treatment, in which abnormalities in p62 and COX protein levels were identified (Fig. 1B-E), revealed an improvement of those protein levels two months after treatment (Fig. 5 C-D). These results suggest that *DNM2* transcript reduction could have an impact on other proteins implicated in CNM pathology.

**Fig. 5:**
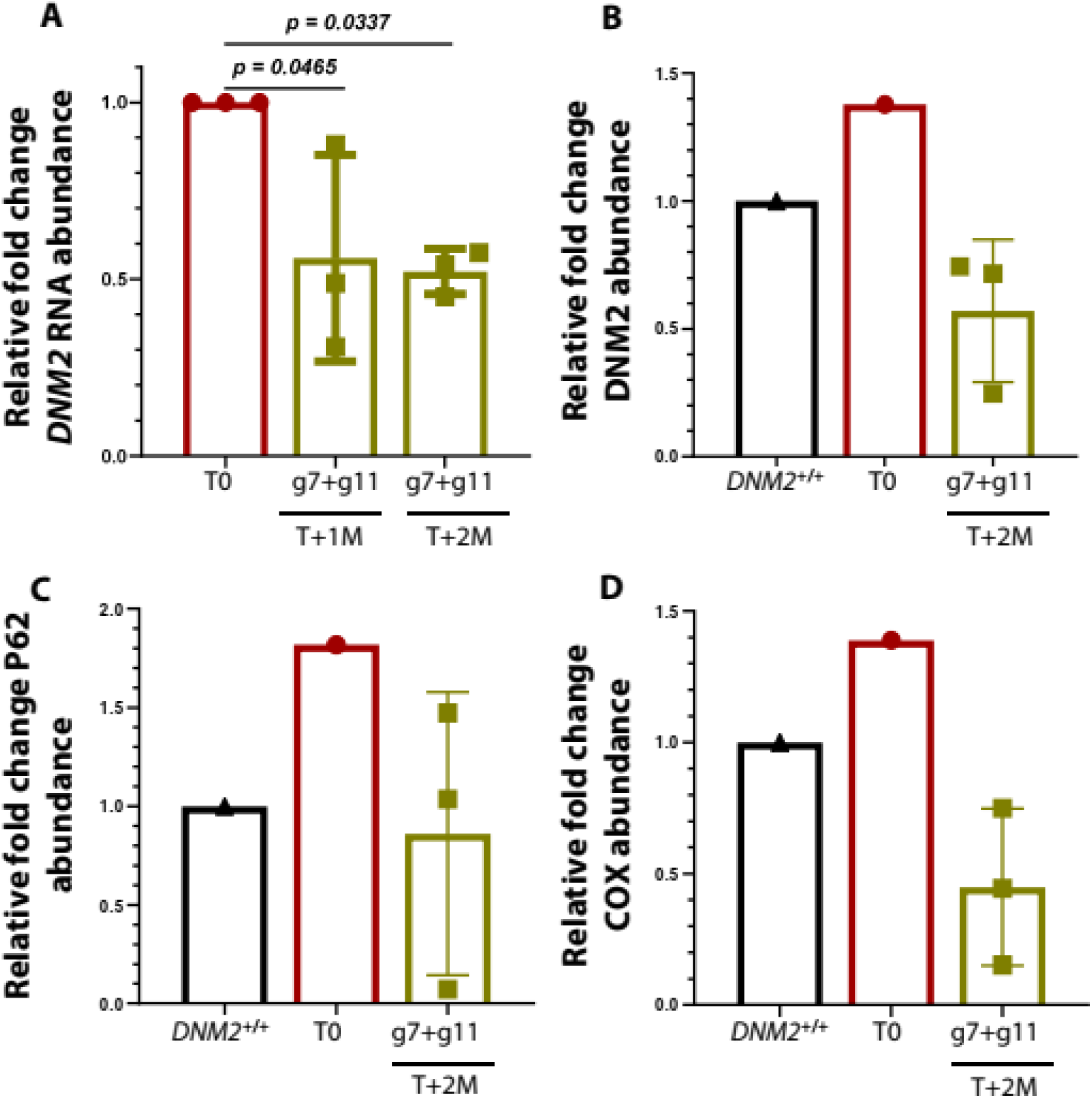
Treatment-induced reduction of DNM2 expression and restoration of autophagy marker levels in dogs. (**A**) RT-qPCR analysis of *DNM2* transcript levels in muscle biopsies from three dogs injected with the AAV-Cas13X.1 and the two gRNAs (AAV-gRNA7 and AAV-gRNA11) at a dose of 3.5×10^12^ vg. Each point represents one dog. Transcript levels were normalized to *GAPDH* and *RPLS19* expression and are expressed as relative fold change abundance relative to baseline (T0) values. (**B**-**D**) Densitometric quantification of WB bands for (B) DNM2, (C) P62 and (D) Cytochrome C oxidase (COX) in treated dogs. Each point corresponds to one injected dog normalized to values of *DNM2^+/+^* control. Data are shown as means ± SEM; One-way ANOVA with Tukey’s test.

### *In vivo* injection does not induce major adverse effects

No local or systemic clinical sign of toxicity was observed in any of the three treated dogs after injections. To assess the safety of the CRISPR/Cas13X.1-mediated *DNM2* reduction approach, we evaluated potential histological, immunological, and transcriptomic effects in treated *DNM2^R465W^*^/+^ dogs. A comprehensive assessment of the fiber morphology, accumulation/infiltration, oxidative activity and fiber type was conducted on the cryosectioned biopsies (Fig. S5 A) by Hematoxylin and Eosin (H&E), Cytochrome C Oxidase (COX), Nicotinamide adenine dinucleotide phosphate diaphorase (NADPH), and myofibrillar ATPase at pH 4.35. Comparative analyses of muscle samples collected before and after treatment revealed no significant overall changes. However, a moderate change was observed in COX staining, characterized by the appearance of “uniformly”-stained fibers (u) and a reduction of fibers unstained (empty, e) fibers (Fig. S5 B). Additionally, a slight shift in fiber type was noted, with a decrease in Type I fibers and increase in Type II (Fig. S5 C).

A key objective of this approach was to assess safety parameters: whether an immune response would be triggered by the CRISPR tool injection or off-target transcripts affected. To this end, we performed a WB analysis to detect serum antibodies against the Cas13X.1 protein in injected dogs. We used proteins lysates from cells expressing both Cas13X.1 and mCherry proteins (using a T2A sequence). The blotted membranes were probed using sera from injected dogs (before and after treatment), as a primary antibody to detect any humoral immune response against the Cas13X.1 protein. The results showed no immunoreactivity corresponding to the molecular weight of Cas13X.1, neither before nor after treatment (Fig. 6A). Given the absence of a commercial anti-Cas13X.1 antibody, we used an anti-mCherry antibody to confirm the presence and detectability of the Cas13X.1 proteins on the blot (Fig. 6B). As a positive control for immune response detection, a serum sample from a Cas9-injected dog and an anti-Cas9 antibody were used, validating the assay’s sensitivity.

**Fig. 6:**
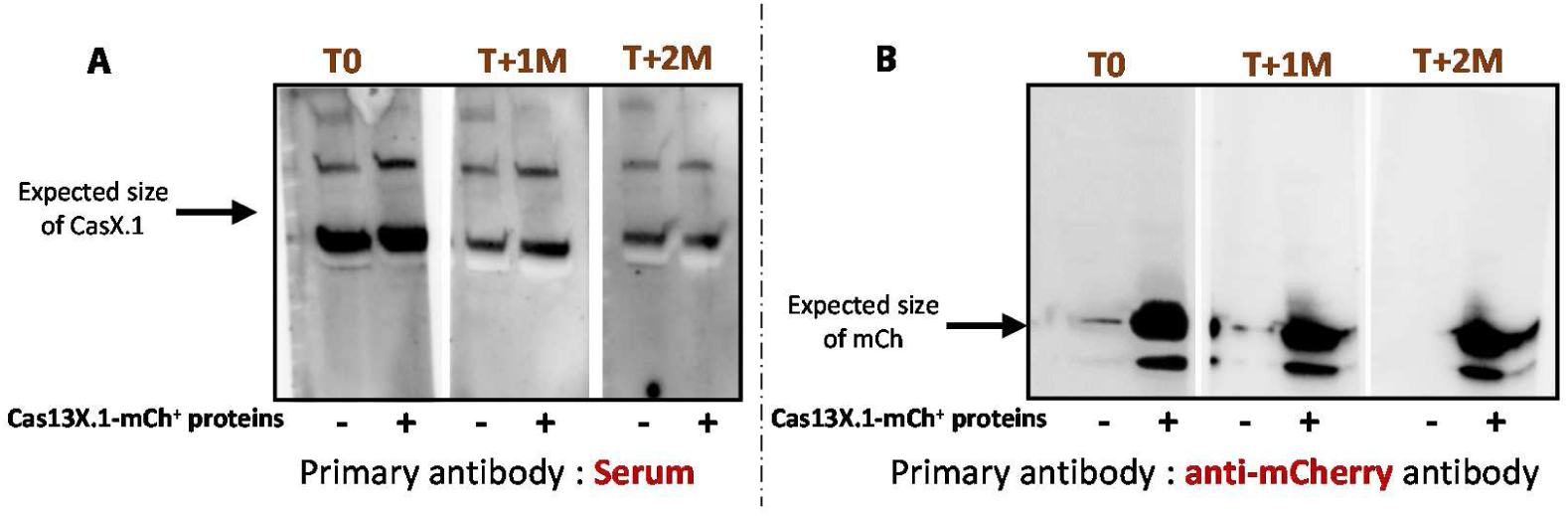
Absence of anti-Cas13X.1 antibodies in treated dogs. Western blot analysis was performed using protein lysates from transfected and FACS-sorted cells expressing Cas13X.1-mCherry (right lane) or control cells lacking expression (left lane). **(A)** Membranes were probed with sera collected from a dog injected with the CRISPR-Cas13X.1 system at baseline (T0), 1 month (T+1M), and 2 months (T+2M) post-injection. (**B**) As a positive control for the presence of the Cas13X.1 protein, membranes were probed with an anti-mCherry antibody.

To evaluate the global impact of the treatment on potential transcriptomic off-target effects, RNA-seq analysis was performed on muscle biopsies from dogs at T+1M. Principal component analysis and differential gene expression analysis revealed no major clustering or deviations between injected and control samples, indicating that global transcriptome integrity was preserved (Fig. S6 A-C). In addition, RT-qPCR analysis of candidate off-targets transcripts sharing partial sequence homology with the gRNAs, *RBBP5*, *CFLAR* and *TSPAN*, revealed no significant decreased expression at T+1M and T+2M, in contrast to the observed downregulation of *DNM2* transcript (Fig. S6 D-G).

Altogether, these results support the safety and tolerability of AAV9-mediated CRISPR/Cas13X.1 delivery *in vivo*. No evidence of muscle toxicity, off-target effects, or immune activation was detected, positioning this approach as a promising therapeutic strategy for CNM.

## DISCUSSION

This study demonstrates the efficacy of the CRISPR-Cas13 system in reducing *DNM2* mRNA expression both *in vitro* and *in vivo* in the *DNM2^R465W^*^/+^ canine model of *DNM2*-related CNM. In cultured myoblasts, we observed a significant decrease of *DNM2* mRNA levels, accompanied by a reduction in protein expression. This led to a reversal of the characteristic cellular alterations, including GTPase activity and the protein level of autophagy-related markers, such as P62. These results were further supported by *in vivo* proof-of-concept experiments, in which intramuscular injections in three *DNM2^R465W^*^/+^ dogs confirmed these results.

The DNM2 protein is a GTPase involved in the fission of endocytic vesicles during clathrin-mediated endocytosis [49], cytoskeleton interaction [15] and autophagy regulation [50]. The R465W mutation is thought to confer a gain of function to the DNM2 protein. Consistent with this hypothesis, we observed an increase in GTPase activity in myoblasts derived from *DNM2^R465W^*^/+^ dogs, as indicated by elevated inorganic phosphate (Pi) release resulting from the hydrolysis of GTP to GDP. These findings align with those of Wang et al. [43], who similarly reported enhanced GTPase activity and an increased oligomer stabilization in cells expressing the *DNM2^R465W^*^/+^ mutation, further supported by Kenniston’s studies [42].

A key functional consequence of altered DNM2 activity is its effect on endocytosis. We observed an increased transferrin uptake in *DNM2^R465W^*^/+^ cells, consistent with observations in murine myoblasts reported by Rabai et al. [33]. However, conflicting results have been reported in the literature. For instance, Bitoun et al. demonstrated by IF that several mutations, including R465W, led to reduced transferrin internalization in COS-7 cells transfected with *DNM2* constructs [51]. Similarly, Trochet et al. observed decreased transferrin uptake by IF in fibroblasts from patients carrying this mutation [52]. Transferrin endocytosis appears to be affected differently depending on the cellular model used. These discrepancies may be due to the type of cells used, such as myoblasts, fibroblasts and COS cells, which originate from a monkey kidney-derived cell line. In addition, the assessment methods differ: our study uses active uptake measurement by quantitative flow cytometry (FACS), whereas others relied on IF-based labeling of associated proteins, a more qualitative method.

We also identified autophagy defects in *DNM2^R465W^*^/+^ cells *in vitro*, characterized by altered expression of the autophagy markers P62 and LC3. LC3 exists in two forms: LC3I (the cytosolic form) and LC3II (the lipidated form). LC3II is associated with the autophagosomal membrane and is considered an indicator of autophagosome formation [53, 54]. p62 (SQSTM1), a key receptor protein in autophagy, plays a central role in the recognition and degradation of dysfunctional proteins and organelles via the autophagosome [55]. The release of COX suggests a potential activation of degradation mechanisms in damaged mitochondria, although it is not directly involved in autophagy [56, 57]. Notably, we observed an increase in LC3-II protein levels, consistent with previous findings [33]. However, the unexpected decrease in p62 levels would typically suggest enhanced autophagic clearance. The simultaneous accumulation of LC3-II and depletion of p62 appears in apparent contradiction, strongly suggesting a profound impairment in autophagic flux rather than a simple induction or blockage of the pathway. This reduction, observed *in vitro*, is particularly intriguing given that p62 protein levels are increased in the muscles of affected dogs. Interestingly, Rabai et al. reported no changes in p62 levels *in vitro* in murine myoblasts, regardless of treatment with an autophagy inhibitor. By contrast, Durieux et al. observed a concurrent increase in both LC3-II and p62 in the liver of knock-in KI-*Dnm2^R465W^* mice, as well as in fibroblasts derived from these animals [40]. Of note, while LC3-II quantification was consistently performed using WB throughout the study, p62 levels were assessed by WB *in vivo* but measured by IF in fibroblasts. Methodological differences may partly account for the results variability. Taken together, these data indicate that the *DNM2^R465W^*^/+^ mutation disrupts the autophagic process, but its precise effect on flux is highly context-dependent. Future studies employing direct flux assays are necessary to fully elucidate the underlying mechanisms in *DNM2*-CNM.

The CRISPR-Cas13 technology is relatively recent and rapidly evolving, with efforts underway to identify smaller and more efficient variants [39, 58]. We specifically chose the Cas13X.1 variant due to its compact size, ideal for AAV vector packaging, and its demonstrated efficacy in reducing *DNM2* expression in our *in vitro* models. To allow for combinatorial testing of different guide RNAs *in vivo* if needed, we packaged the Cas13 nuclease and the gRNAs into separate AAV vectors for intramuscular (IM) injection. We conducted IM injections using the Cas13X.1, and the most effective gRNAs identified *in vitro*. Our *in vivo* experiments confirmed the efficacy of this strategy, demonstrating a significant reduction of both *DNM2* mRNA and protein levels, particularly at T+2M post-injection. The muscle bundles injected were entirely harvested at T+1M and T+2M, which limits our analysis to a short-term therapeutic window for this proof-of-concept experiment. Despite this robust molecular correction and the normalization of biochemical markers like COX and p62 proteins, we did not observe significant improvements in muscle histology. The injected dogs were five years old at the time of treatment and their muscles had therefore been diseased for a long period of time (onset of histological signs: two months of age). The reversibility of the histological phenotype at this stage can therefore be questioned. Future systemic injections will be essential to evaluate long-term functional benefits driven by the treatment. Whereas affected BF muscles of *DNM2^R465W^*^/+^ dogs exhibited a slight increase in Type I fibers, in treated BF muscles we observed a partial change, with type I fibers decreasing to levels comparable to those of Type II fibers, however these changes were not significant. Long-term treatment, particularly with systemic administration in younger animals, will be essential to fully assess therapeutic outcomes, or even prevent them. This hypothesis is supported by work in *Dnm2^R465W/+^* mice, where early intervention with *DNM2*-targeting shRNA prevented disease onset and corrected histological abnormalities [13].

Translating the success of local administrations into a systemic therapy will require vector optimization and long-term safety follow-up. In this study, we used AAV9 for IM injections; however engineered myotropic AAVs, such as Myo-AAV, provide improved skeletal muscle specificity and reduced liver transduction. Myo-AAVs have shown greater efficacy in muscle transduction, which could be critical for durable and targeted therapies in the treatment of myopathies [59, 60]. Additionally, a more thorough evaluation of the immune response induced by AAV-CRISPR/Cas13 systemic injections will be necessary to ensure the long-term safety of this therapy. So far, WB analyses performed with sera from the three injected dogs have shown no humoral response against Cas13X.1, suggesting good tolerability. This is encouraging for translational prospects, especially given that Cas13X.1 is currently being tested in clinical trials for the treatment of age-related macular degeneration, where it is delivered via one single AAV vector to partially down the expression of the target, the VEGF (NCT06031727). These developments underscore the therapeutic potential and clinical relevance of Cas13-based RNA targeting for a range of genetic diseases, including *DNM2*-related CNM.

Our results support a growing consensus that a ∼50% reduction of DNM2 expression is sufficient to mitigate key pathological features of CNM, aligning with previous findings from studies using shRNA or antisense oligonucleotides [13, 22]. However, the use of CRISPR-Cas13X.1 in our study offers advantages, particularly in terms of potentially enhanced specificity by targeting *DNM2* mRNA without interfering with other transcripts, thus reducing the risk of off-target effects in a whole-body delivery approach [36, 37]. The clinical viability of this approach is underscored by the ongoing first-in-human trials of Cas13X.1-mediated VEGF knockdown in age-related macular degeneration (NCT06031727). Our molecular analyses demonstrated a clear reduction in *DNM2* transcript and protein levels in treated muscle, with no apparent off-target effects among the genes assessed. Nonetheless, further transcriptome-wide studies, on the longer term will be necessary to confirm the absence of off-target effects and ensure the safety of this potential therapeutic approach.

A key strength of our study lies in the use of a canine model, which more closely recapitulates the human disease phenotype and progression. The *DNM2^R465W^*^/+^ canine model exhibits muscle pathology and clinical progression that closely mirror those of affected patients, thereby enhancing the translational relevance of our findings. The ethical and practical constraints on animal availability imposed limitations on our proof-of-concept experiments, which were restricted to three dogs. This number was sufficient to highlight the reduction in DNM2, which demonstrates its robustness. However, it is possible that this number was too small to show significant differences on other parameters, such as histological improvement.

Despite these limitations, our combined *in vitro* and *in vivo* data provide strong support for the adaptation of this technology in future clinical trials. In conclusion, this study provides a critical preclinical proof-of-concept for AAV-mediated CRISPR-Cas13 therapy for *DNM2*-related centronuclear myopathy. The treatment was well tolerated and there was no evidence of muscle toxicity or immune response against the Cas13 protein. We observed a reproducible ∼50% reduction in *DNM2* mRNA levels in injected muscle samples, accompanied by a corresponding decrease in protein expression. Furthermore, the partial reversal of biomarkers, such as p62 and COX, suggests that defects in autophagy and mitochondrial homeostasis could be amenable to correction through systemic treatment. These findings establish a proof of principle for the continued development of RNA-targeting CRISPR therapies, offering a promising new avenue for treating CNM and other dominant genetic disorders for which no curative therapies currently exist.

## MATERIALS AND METHODS

### Study design

All the *in vivo* procedures were performed in accordance with the EU Directive 2010/63/EU for animal experiments and were approved by the Ethical committee of EnvA, ANSES and UPEC (agreement number #16) and by the French research ministry (APAFiS nbr #13015-2018010910531134 v3, #37794-2022062411148501 v3 and #43589-2023051700273252_v5).

The sample size for the *DNM2^R465W/+^* dog studies was determined based on the in vitro range of *DNM2* mRNA reduction and its variability. For ethical reasons, we used the dogs that we had already obtained to characterize the disease rather than producing new litters. Therefore, intramuscular injections were performed in adult dogs, which is a highly-relevant translational condition (mimicking adult patients). The injection was performed on muscle bundles as a proof of principle of CRISPR/Cas13 effectiveness in reducing *DNM2* transcripts. Sample sizes for each experiment are included in the figure legend. Information on sample collection, treatment and processing has been included in the results section and other materials and methods sections. To increase our number of primary myoblasts from healthy dogs, we punctually added data from healthy Golden Retriever dog myoblasts available in the laboratory as specified in the figure legends. Primary myoblasts were obtained from biopsies taken from dogs at 24 months of age. Immortalized myoblasts were obtained from these primary myoblasts according to the process described in another section of the materials and methods. No outliers were excluded from the study.

### Biopsies

Biopsies were used for cell culture, for histological analysis and for RNA-Seq. They were surgically taken from the *biceps femoris, triceps brachii* or *tibialis cranialis* muscles by a surgical operation. The dogs were anesthetized using an intravenous induction of propofol (6.5 mg/kg), and anesthesia was maintained with isoflurane in 100% O2. Analgesia was ensured by an intravenous injection of morphine (0.1 mg/kg). Continuous monitoring of ECG, SpO2, ETCO2 and rectal temperature was set up, and an automatic ventilator was used to maintain normocapnia. Surgical biopsies were taken and immediately vertically mounted on a piece of cork, using tragacanth gum, and snap-frozen in isopentane cooled in liquid nitrogen except for the parts of the biopsies used to obtain muscle cells. Samples were stored at −80°C prior to sectioning and performing histological analysis.

### Cryosection histological analysis

#### Muscle morphology

Transverse sections (10 µm) from cryopreserved muscle biopsies were stained with H&E, ATPase pH 4.35, modified Gomori trichrome, NADH tetrazolium reductase (NADH-TR) and cytochrome c oxidase (COX), and assessed for fiber morphology, accumulation/infiltrations, and oxidative activity. Quantitative analyses were performed on H&E staining. Entire muscle sections were analyzed using Visilog 7.0 software (Noesis). The Visilog 7.0 software was used for quantification of COX and NADH-TR muscle abnormalities in pre and post-treated muscles.

#### Muscle immunostaining

Seven µm slices cryostat sections were fixed with acetone/methanol for 10 min at −20°C. Nonspecific sites were blocked with 3% - Bovine Serum Albumin in Phosphate-buffered saline (PBS) 1 h, room temperature. Tissue slices were stained with the primary antibody in PBS + 3% BSA (overnight, 4°C), then with anti-mouse Alexa Fluor 647 in PBS+ 3% BSA (1 h, RT) and an anti-rabbit Rho. Nuclear staining was performed with DAPI (4’,6-diamidino-2-phénylindole), Sigma-Aldrich (300 nM). Primary antibody: mouse anti-p62 (Abcam ab56416).

### Canine myoblasts isolation and culture

*Biceps femoris* muscle biopsies were performed to obtain *in vitro* myoblasts culture for the characterization of the phenotype. The biopsy was cut into fragments of about 1 to 2 mm and then rinsed with PBS + 2% Fetal Bovine Serum (FBS) (South America, SV30160.03) to remove blood and toxins from damaged fibers. After that, the biopsy was digested by type II collagenase (Worthinton) for 2 hours at 37°C. The mixture was passed through an 18G needle and then through 100μm and 40μm strainers). Cells were then centrifuged at 1600 rpm for 5 minutes and washed with PBS + 2% FBS. After a second centrifugation, cells were cultured with custom MCDB120 medium (HyClone Cytiva, RRVA220814) supplemented with FBS at 20%, gentamycin at 25 μg/ml, dexamethasone and FGF-ß at 10 ng/ml. Cells were cultured in a humidified incubator at 37 °C with 20 % O2 and 5 % CO2. Cells were analyzed by the expression of Desmin and CD56 as myogenic markers. Myotube formation was induced by culturing myoblasts in DMEM supplemented with 2 % horse serum and 1% Penicillin/streptomycin for 3 to 7 days. Cells were labeled and analyzed by flow cytometry (FCM) using FACS (BD Accuri C6 PLUS Ref.123720). FACS allowed us to determine the percentage of myogenic cells by detecting the expression of both myogenic markers CD56/NCAM (Santa-Cruz sc-1507) and clone D33 (Dako M0760) among our cell population. Cell labeling was done in 96 well U-shaped plates with at least 100,000 cells/well.

### Myoblast’s differentiation and fusion: MHC labeling and primary fusion index

Cells were cultured in a culture chamber (Labtek) suitable for microscopy and cell differentiation was induced using differentiation medium, composed of DMEM supplemented with 2% horse serum and penicillin/streptomycin. The medium was changed after two days until seven days of differentiation. After 7 days, we fixed our cells with methanol ten minutes at −20°C.

For the IF experiments, cells were blocked with a 3% BSA-PBS solution for 1 hour and incubated at room temperature (RT) with primary antibody against Myosin Heavy Chain (MF20-c DSHB) and then incubated with proper secondary antibody for 30 min (stained with Alexa Fluor 488nm AbCam). Cells were washed three times between both incubations. Nuclei were stained with DAPI for 5min. Negative control samples: secondary antibody received equivalent amount of Alexa Fluor 488- and PE-labeled isotype-matched antibodies while unlabeled cells control was untreated.

### Cell immortalization process

To overcome the problem of limited cells lifespan (also the limited number of biopsies from *DNM2^R465W/+^* dogs), we immortalized cells. *DNM2^R465W/^*^+^ and *DNM2^+/^*^+^ myoblasts were transduced with the lentiviral EF1-hTERT-hCDK4-R-Blasticin vector containing genes for immortalization (hTERT for Human telomerase reverse transcriptase and CDK4 for cyclin-dependent kinase 4). HEK 293T cells cultured in 10 cm petri dishes were transfected (CaPO_4_ transfection mediated by CaCl_2_ and HBS) with 5.2 µg VSV-G plasmid, 9.7 µg PAX2 plasmid, 5.2 µg Tat plasmid and 10 µg pRRL.EF1.hTERT2AhCDK4.2AblastR plasmid. Viral vector particles-containing culture medium was harvested at 24 and 48 hours post transfection. Viral vectors were concentrated by centrifugation with an SW28 rotor (Beckman Coulter) 25,000 rpm, 120 min, 4 °C and resuspended in 90 µl of PBS. Myoblasts (at passage 2) seeded in a six-well plate were transduced with 3µl of viral vector suspension in 300µl of medium. 48h later selection antibiotic was added to cells for transduced cell selection. To select the myogenic immortalized cells, a N-CAM (CD56) Santa-Cruz labeling was performed. The ARIA FACS-sorter was used to sort CD56-positive cells which were then amplified in culture.

### DNM2, p62, LC3 and Desmin labeling on *in vitro* cells

For the IF experiments, cells were blocked with a 3% BSA-PBS solution for 1 hour and incubated at 4°C overnight with primary antibody against: DNM2 anti-rabbit recombinant monoclonal Dynamin 2 antibody (Abcam 151555) diluted 1/100; p62 anti-p62/SQSTM1 (Abcam ab56416) diluted 1/100; LC3I and II; MAP1LC3A/B (N-Terminal) (AHP2167 Bio-Rad) diluted 1/100. Desmin anti-Desmin clone D33 (Dako M0760) diluted 1/100. Cells were then incubated with appropriate secondary antibody for 45 minutes (stained with Alexa Fluor 488nm AbCam, Alexa Fluor 647nm AbCam or Rho 550nm AbCam). Cells were washed with PBS three times between both incubations. Nuclei were stained with DAPI for 5 minutes.

### Measure of GTPase activity and protein analysis on *in vitro* cells

GTPase activity was measured using Sigma-Aldrich “ATPase/GTPase Activity Assay Kit” (MAK113). All samples and standards were analyzed in duplicate. Cells were placed in a 12-well plate with 20000 cells/well. The following day, they were incubated for 30 min with either: medium + 0.2% DMSO or medium + Dynasore 120µ M. The reaction mix was prepared according to the Sigma kit protocol: 20µ L Assay Buffer + 4 mM GTP at 10µ L/well (except for the range) i.e. 30µ L/well and incubated for 30minutes at RT. The absorbance was read using the BioTek SYNERGYLX plate reader at 620nm. Results were normalized by protein quantity per well. After optical density reading, proteins were extracted by RIPA lysis buffer (5M NaCL, 0,5M EDTA at pH 8, 1M Tris at pH 8, NP-40 IGEPAL CA-630, 10% deoxycholate sodium and 10% SDS with 10µ L of anti-protease per mL of buffer) and quantified by Pierce BCA Protein Assay Kit (Thermo Scientific).

### Transferrin uptake for endocytosis studies on *in vitro* cells

Cells were cultured in a 12-well plate (65000/well). The next day, cells were starved with IMDM containing 0.5% BSA for 30minutes. They were then incubated in the same medium containing 20µg/mL Alexa fluor 647 fluorescently-labeled CY5-transferrin (Jackson ImmunoResearch Laboratories), for 10 minutes on ice. Endocytosis is initiated by incubating cells at 37°C for 20 minutes. Cells were washed with 0.02% sodium azide for 3 minutes to remove transferrin remaining on the cell surface. Cells were then detached with trypsin, fixed with 2% PFA for 15minutes at 4°C and resuspended in PBS. Cells were analyzed by FACS Accuri and the mean CY5 intensity measured with FACS software BD Sampler Plus.

### Plasmids

Plasmids for Cas13 proteins and gRNA expression were obtained from Addgene. The LwCas13a-msfGFP-2A-Blast plasmid (Addgene #91924) was used for LwCas13 expression under the EF1α core promoter. The EF1α-CasRx-2A-EGFP plasmid (Addgene #109049) was used for Cas-Rx expression. The CMV-Cas13X.1-SV40pA_U6-BbsI-DR_CMV-mCherry-BGHpA plasmid (Addgene #171379) was used to express Cas13X.1 protein. For gRNA cloning, the LwaCas13a guide expression backbone with the U6 promoter (Addgene #91906) and the RfxCas13d pre-gRNA cloning backbone plasmid (Addgene #109054) were utilized.

### gRNA design and cloning

gRNA design was performed using the Cas13 design page from Sanjana Lab, and RNA secondary structure was analyzed using the RNAfold webserver. The primers for gRNAs (synthesized by Eurofins Genomics) were annealed at 95°C and ligated into linearized plasmids (cut with BpiI/BbsI) using T4 DNA ligase (NEB) (Table S.1A). Plasmids were confirmed by sequencing prior to AAV packaging.

### Cell transfection

Transient transfection of *DNM2^R465W^*^/+^ cells was performed to test the efficiency of the CRISPR/Cas13 tool. For this transfection, when cells reached at least 60% confluence, plasmids were transfected using Lipofectamine 3000 (Thermo Fisher Scientific), according to the manufacturer’s protocol. A dual plasmid system was used for CRISPR/Cas13 expression, with one or two plasmids encoding gRNA under the hU6 promoter, and another plasmid for the Cas13 protein (EF1a-CasX.1-mCherry). Cells were sorted 48 hours after transfection according to mCherry (reporter gene) expression using the BD Aria Fusion sorter. Cells were used for GTPase measurement, or frozen for protein and/or RNA extraction for further analysis.

### RNA isolation and Reverse Transcription-quantitative Polymerase Chain Reaction

RNA was isolated from canine cells or frozen muscle biopsy sections. Total RNA was extracted using QIAzol. Extraction quality was quantified using the Thermo Fisher Scientific^TM^ NanoDrop^TM^. Reverse Transcription (RT) was performed using Thermo Fisher Scientific “Maxima first strand cDNA synthesis kit for RT-PCR”. Quantitative real time PCR (qPCR) was realized with SYBR Green dye (Thermo-Fisher Ref. K0252) using a QuantStudio 3 Applied Biosystems. RT-qPCR data was analyzed using 2 delta delta Ct method (2ΔΔct) to quantify fold change between controls and tested samples. Primer sequences are listed in Table S.1B

### Western-Blot

Proteins were extracted using RIPA buffer containing a cocktail of protease inhibitors (P2714 Sigma-Aldrich). WB were performed on proteins extracted from muscle biopsy sections or immortalized and primary cells. Protein concentration were assayed using the BCA protein assay kit (Bio-Rad). Between 9 and 20 µg of protein are denatured at 95°C for 5 min. Samples were loaded onto Mini-PROTEAN TGX Stain-Free Precast Gels (4-15%) and electrophoresed for 30 minutes at 200 volts. The Stain-Free gels were UV-activated using the Bio-Rad ChemiDoc MP Imaging System to visualize total proteins. Samples were transferred to PVDF membranes using the Invitrogen iBlot 2. Non-specific sites were blocked using PBS containing 5% milk for one hour. Antibodies: DNM2 Rabbit Recombinant Monoclonal Dynamin 2 antibody Abcam 151555 diluted 1/1000. The Anti-p62/SQSTM1 (ab56416 Abcam) and LC3 I and II proteins human MAP1LC3A/B (N-Terminal) (AHP2167 Bio-Rad) are diluted 1:1000. BD Pharmingen™ Purified Mouse Anti-Cytochrome c antibody (556433) is diluted 1:500 in PBS-3%BSA. For WB using sera as primary antibodies, the dilution used is 1/100. All primary antibodies were incubated at 4°C overnight. Membranes are then incubated with peroxidase-conjugated secondary antibodies diluted 1:5000 in PBS-3%BSA for 45 minutes at RT and revealed with an enhanced chemiluminescence (ECL) kit (Thermo Fisher Scientific) using Bio-Rad ChemiDoc MP Imaging System. Protein quantification was then performed using ImageJ software.

### *In vivo* intramuscular injections

The dogs were anesthetized using the same protocol as for muscle biopsies. The targeted muscle was first exposed and injection zones were delineated in distant bundles using non-absorbable sutures (Prolene 4-0). Intramuscular injections. Intramuscular injections were performed between the two markers, using a Hamilton syringe and a 27G needle was used at a rate of 10 injections of 50 µl each, for a total injected volume of 500 µl were performed.

### AAV vector cloning and production

The Cas13X.1 protein cDNA was excised from plasmid #171379 using AgeI and SpeI restriction enzymes, purified, and cloned into the AgeI and NheI sites of the AAV plasmid backbone (Addgene #109049). The resulting plasmid was the ITR-EF1α-NLS-Cas13X.1-NLS-PolyA-ITR.

The AAV/9 vectors obtained for the *in vivo* experiments were manufactured by PackGene Biotech. Viral vectors were purified by iodixanol gradient ultracentrifugation and a step of concentration. Physical particles were quantified by real-time PCR. AAV purity was assayed by SDS-PAGE electrophoresis using Coomassie Blue Staining. AAVs were tested for endotoxin free status. Titers were adjusted to 2×10^13^vg/ml.

### Bioinformatic analysis of RNAseq data

The transcriptome analysis (Total RNA-Seq), including ribosomal RNA depletion, was performed by Integragen (Evry, France), using the Illumina® NovaSeq X Plus (100 million pairs, PE100 sequence). FASTQ files were aligned to the dog reference genome (Ensembl CanFam 3.1 assembly) using the Hisat2 aligner software (Hisat2.2.1) with paired-end parameters [61], and exported into BAM format [62] or SALMON for FPKM values. An average of 100 million reads per sample were obtained, with a mean mapping efficiency of 86%. FeatureCounts was used to obtain the number of aligned reads per gene [63]. Log2 transformed FPKM values of expressed transcripts between control (First injected dog at T0 and dogs at 24 months) and other conditions were plotted with ggplot2 (https://doi.org/10.1007/978-3-319-24277-4).

### Statistical analysis

The data are expressed as means ± SEM. Graph and curves were made using GraphPad Prism software versions 6 and 10. The unpaired Student’s t-test was used to compare two groups. Non-parametric tests, including Kruskal-Wallis or Dunn’s post hoc tests were used to compare multiple groups. One-way ANOVA and Bonferroni or Tukey’s post hoc test were used to compare different groups if the data followed a normal distribution and if the samples analyzed had the same genetic background. P values smaller than 0.05 were considered significant. The number of dogs and myoblasts from different dogs, as well as the tests used for each experiment, are indicated in the figure legends.

## List of Supplementary Materials

Fig. S1 to Fig S6

Tables S1 and S2

## Acknowledgments

We would like to express our sincere gratitude to ZNM – Zusammen Stark! e.V. for their financial support, which has made this project possible. We would also like to express our gratitude to the Association Française contre les Myopathies (AFM-Téléthon), the Pequeños Superhéroes association, the Agence Nationale de la Recherche (Laboratoire d’Excellence Revive), and the Fondation Maladies Rares for their support and funding. We would like to express our gratitude to the IMRB bioinformatics core lab for their assistance with RNA-Seq analysis and to the IMRB cytometry platform for their support with cell sorting.

## Author contributions

Conceptualization: IP Methodology: AC, IB, LT, SB, IP

Performed experiments: AC, IB, NBG, SCJ, FA, TO, IP

Funding acquisition: FR, SB, IP

Writing – original draft: AC, IP

Writing – review & editing: AC, IB, NBG, SCJ, ND, FR, LT, IP

## Competing interests

IP has a patent related to the data reported in this paper (#FR2501167). The other authors declare no conflict of interest.

**Data and materials availability**: all data associated with this study are present in the paper or the supplementary materials.

