## Supplementary figures for "CRISPR/Cas13-mediated Dynamin 2 reduction therapy in a canine model of *DNM2*-related centronuclear myopathy"

SUPPLEMENTARY FIGURES S1 – S6

SUPPLEMENTARY TABLES 1 AND 2

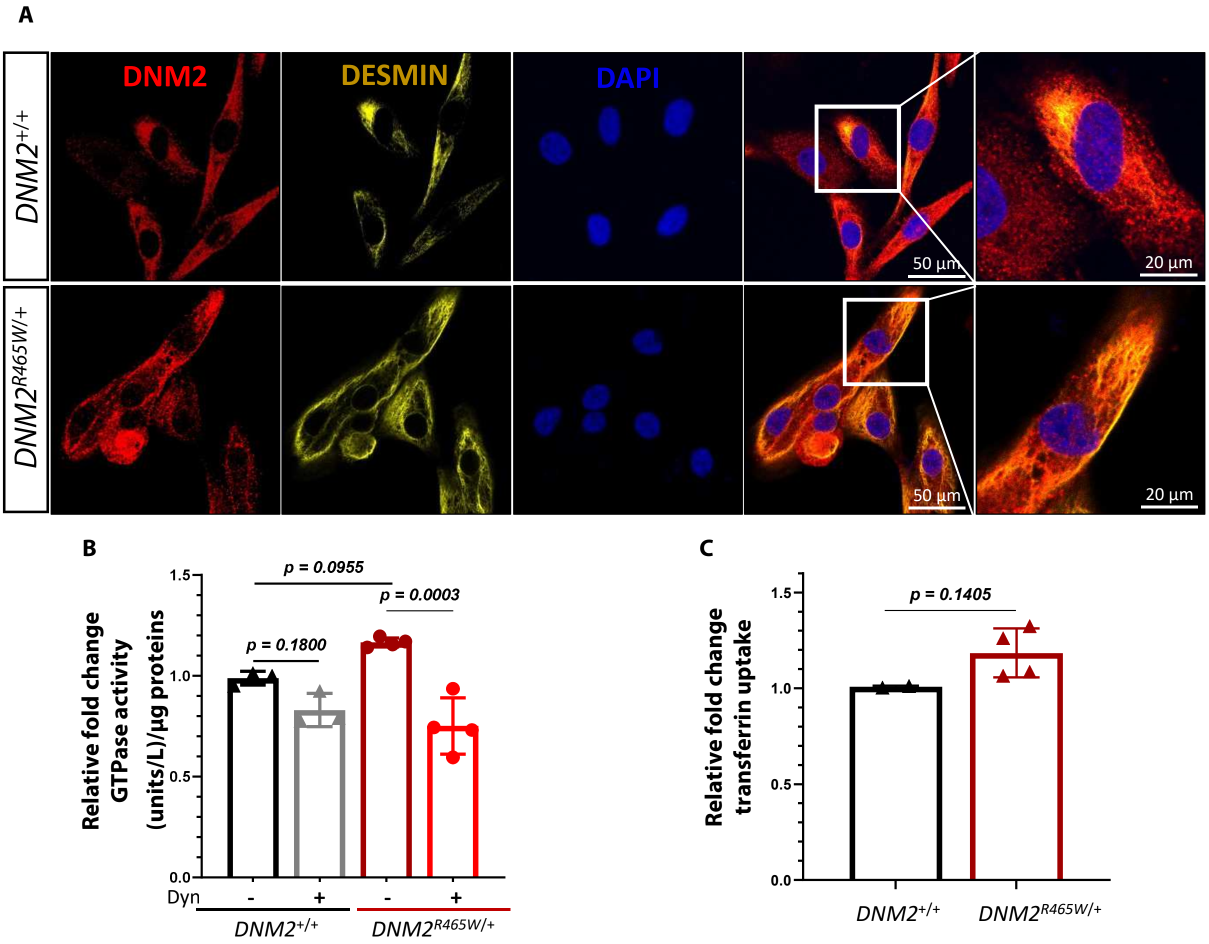


**Fig. S1:** ***DNM2$^{R465W/+}$*** **immortalized myoblasts show alterations in GTPase activity and endocytosis.**
(**A**) Immunofluorescence staining for Dynamin 2 (DNM2) and Desmin (DESMIN) on *DNM2$^{+/+}$* (top panel) or *DNM2$^{R465W/+}$* (bottom panel) cells. Nuclei are counterstained with DAPI. (**B**) GTPase activity in immortalized cells from muscle biopsies of four *DNM2$^{R465W/+}$* dogs, with or without treatment with a Dynamin inhibitor, Dynasore (Dyn); normalized to values of their matched control healthy littermate and two additional *DNM2$^{+/+}$* controls. (**C**) Quantification of Cyanine3-labelled transferrin uptake in immortalized cells from muscle biopsies of four *DNM2$^{R465W/+}$* dogs, normalized to values of their matched control healthy littermate and an additional *DNM2$^{+/+}$* controls. Data are shown as means ± SEM per dogs ; One-way ANOVA with Tukey's post hoc test (B) and Unpaired t test (C).

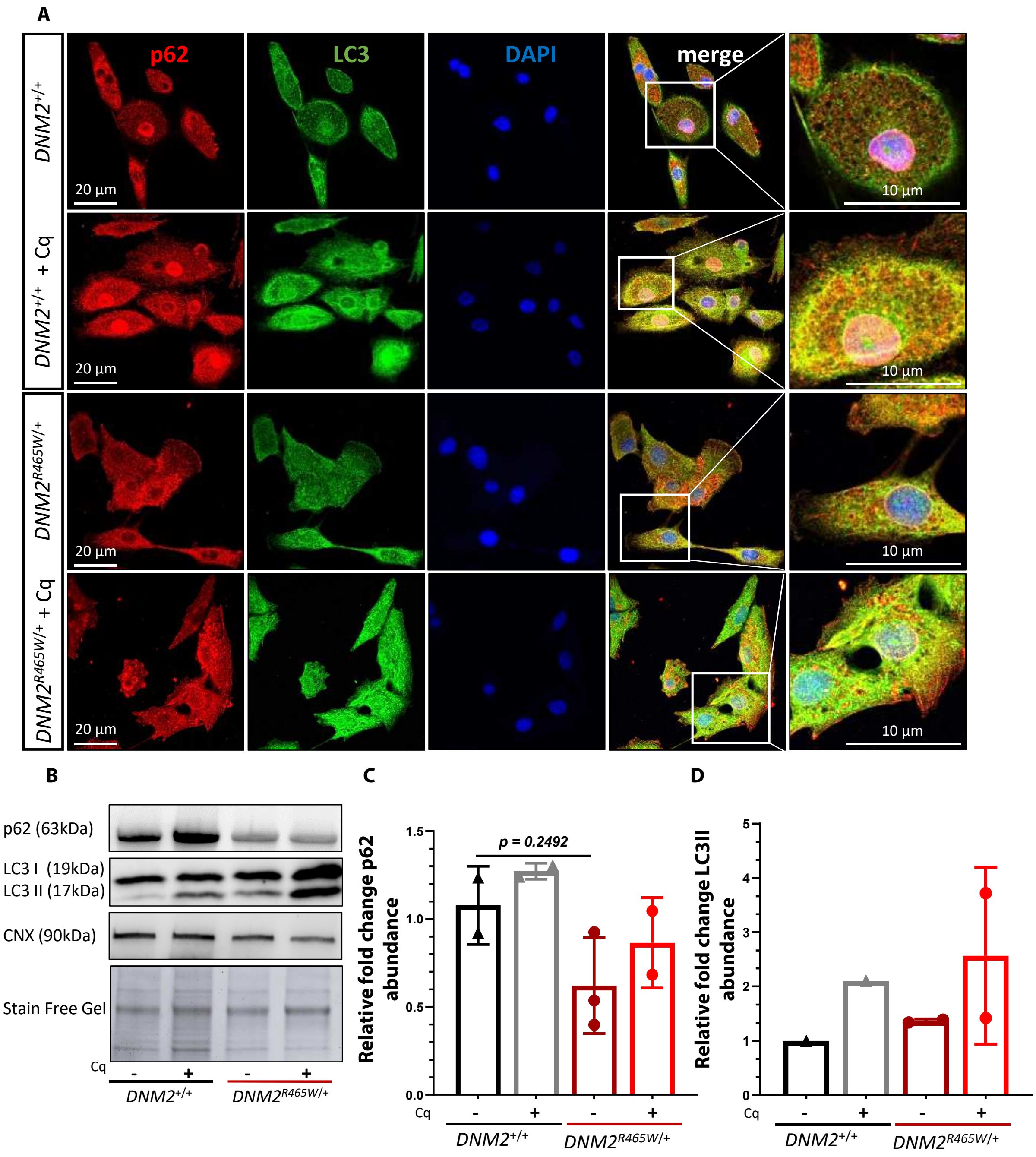


**<u>Fig. S2:</u> *DNM2*$^{R465W/+}$ immortalized cells maintain alterations in autophagy protein expression.**

(**A**) Immunofluorescence staining for P62 and LC3 on *DNM2*$^{+/+}$ (top panel) and *DNM2*$^{R465W/+}$ (bottom panel) immortalized cells, with or without treatment with chloroquine (Cq) which blocks autophagy (50µM for 16 hours). Nuclei are counterstained with DAPI. (**B**) Representative WB performed on immortalized cells, with or without treatment with Cq, staining for P62, LC3 and Calnexin (CNX). (**C**) Quantification of the intensity of P62 marker protein bands in immortalized myoblasts from muscle biopsies of four *DNM2*$^{R465W/+}$ dogs, with or without the addition of Cq; normalized to values of two *DNM2*$^{+/+}$ control dogs. (**D**) Quantification of the intensity of LC3 II marker protein bands in immortalized cells from muscle biopsies of two *DNM2*$^{R465W/+}$ dogs, with or without treatment with Cq; normalized to values of *DNM2*$^{+/+}$ control. Data are shown as means ± SEM per dogs ; One-way ANOVA with Tukey's test.

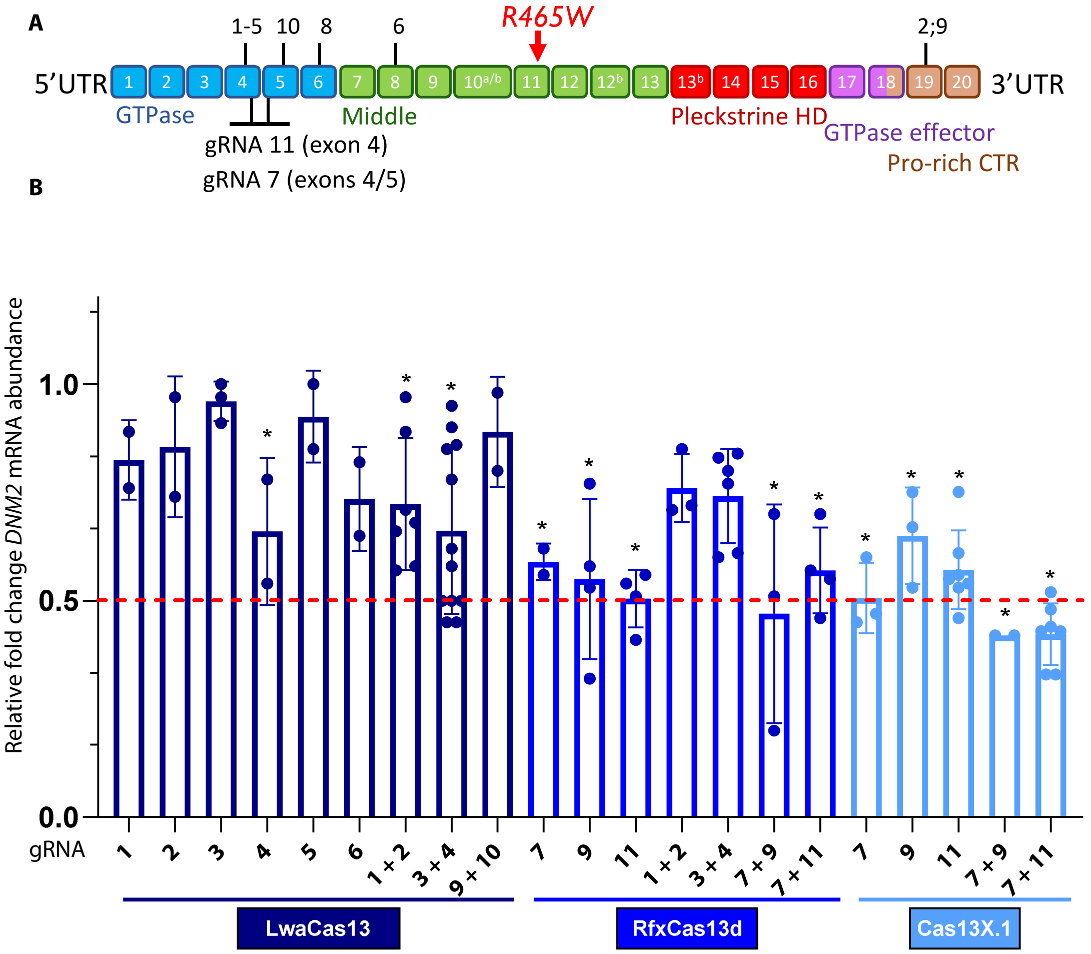


**Fig. S3 :** ***In vitro* knockdown of *DNM2* mRNA by CRISPR/Cas13**

(**A**) Schematic representation of the DNM2 protein domains and position of the R465W mutation (red arrow). Targeted sequences of the two selected gRNAs and positions (black lines) of other gRNAs tested in the CRISPR-Cas13 system on *DNM2*$^{R465W/+}$ cells. (**B**) RT-qPCRs performed on RNA extracted from transfected and FACS-sorted cells. Each point corresponds to an independent experiment (for each point, 3 qPCR triplicates). *GAPDH* and *RPLS19* used as housekeeping genes and results are expressed as the normalized (to the Cas13 with the empty vector) relative fold change *DNM2* mRNA abundance. The red line corresponds to the target of a 50% reduction in *DNM2* mRNA. One-way ANOVA with Tukey's test *$p < 0.05$.

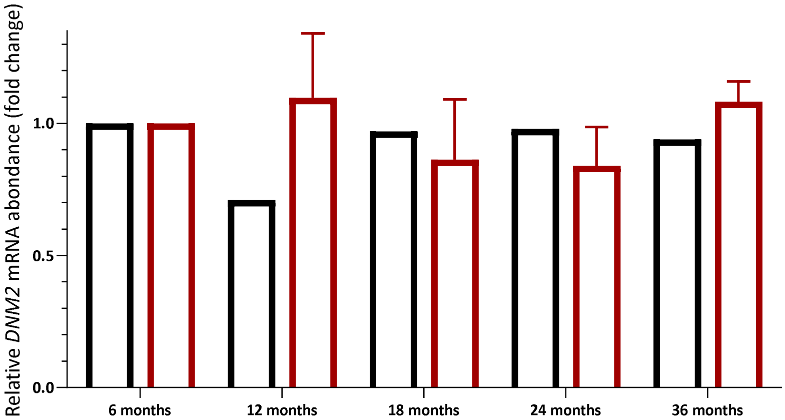


**<u>Fig. S4:</u> Stable *DNM2* RNA levels over time in *DNM2*$^{R465W/+}$ dog muscles.** RT-qPCR analysis of the mRNA content of *DNM2* from the BF muscle of four *DNM2*$^{R465W/+}$ dogs (red) and the healthy *DNM2*$^{+/+}$ dog (black) at 6, 12, 18, 24 and 36 months. The mRNA was referred to *GAPDH* and *RPLS19* housekeeping genes and normalized results are expressed as *DNM2* mRNA relative fold change abundance related to the biopsies taken at 6 months.

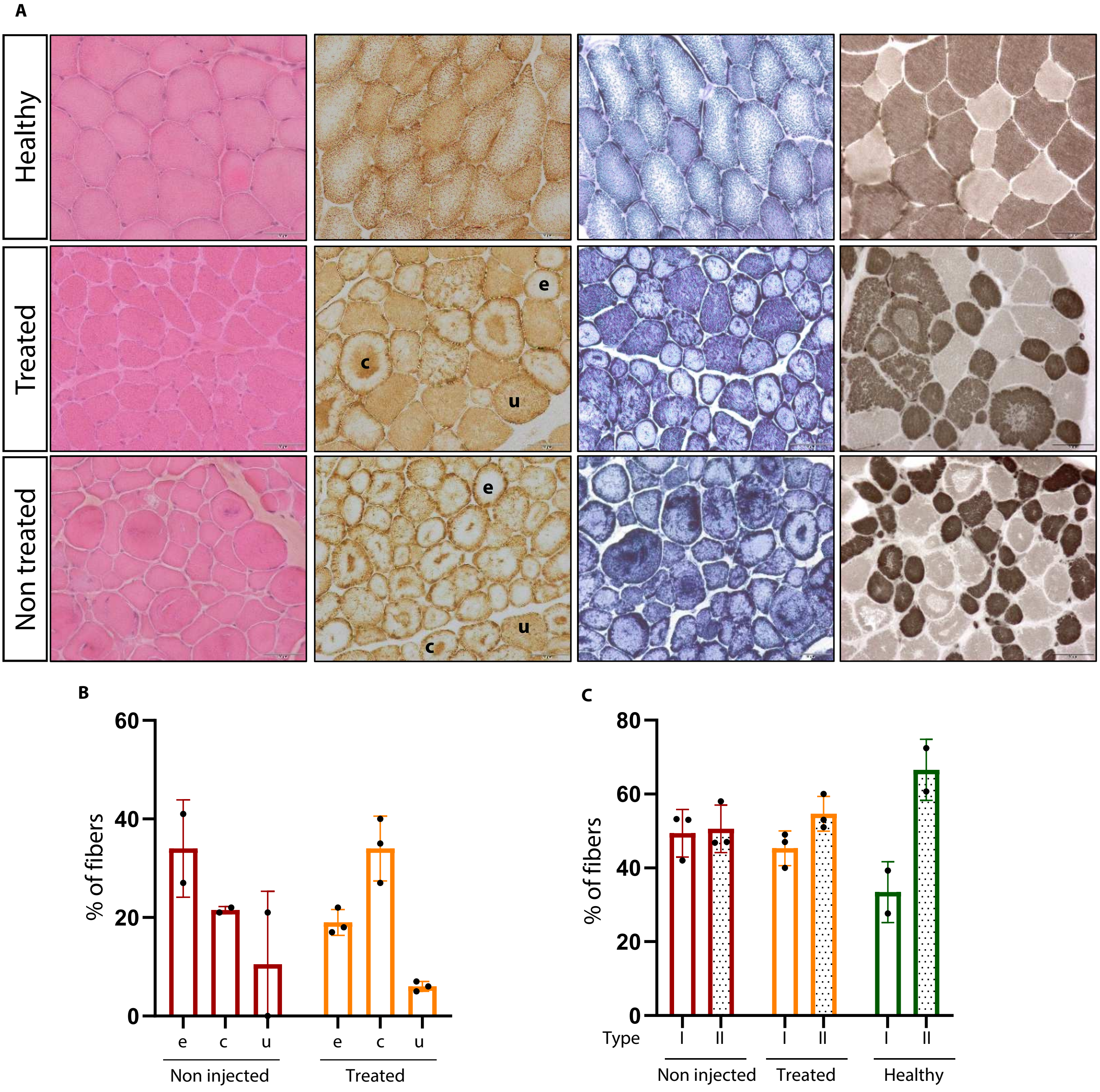


**<u>Fig. S5:</u> Histological and histochemical analysis of injected muscles.**
(**A)** Representative serial cross-cryosections of BF muscle biopsies from a healthy control (top row), a treated dog (middle row), and an untreated dog (bottom row). Columns from left to right show staining for: Hematoxylin and Eosin (H&E), Cytochrome C Oxidase (COX), Nicotinamide adenine dinucleotide phosphate diaphorase (NADPH), and myofibrillar ATPase at pH 4.35. Scale bar: 50µm. (**B)** The COX staining was used to quantify the percentage of "empty" fibers (e), "condensate" (c) and "uniform" stained fibers (u) in non–injected (left, in red) and treated (right, in orange) muscles. (**C)** ATPase staining was used to quantify percentage of Type I (empty columns) and Type II (punctuated columns) fibers. Each point corresponds to a treated dog muscle. The non-injected points correspond to the dog-matched T0 and 24 old-months muscles.

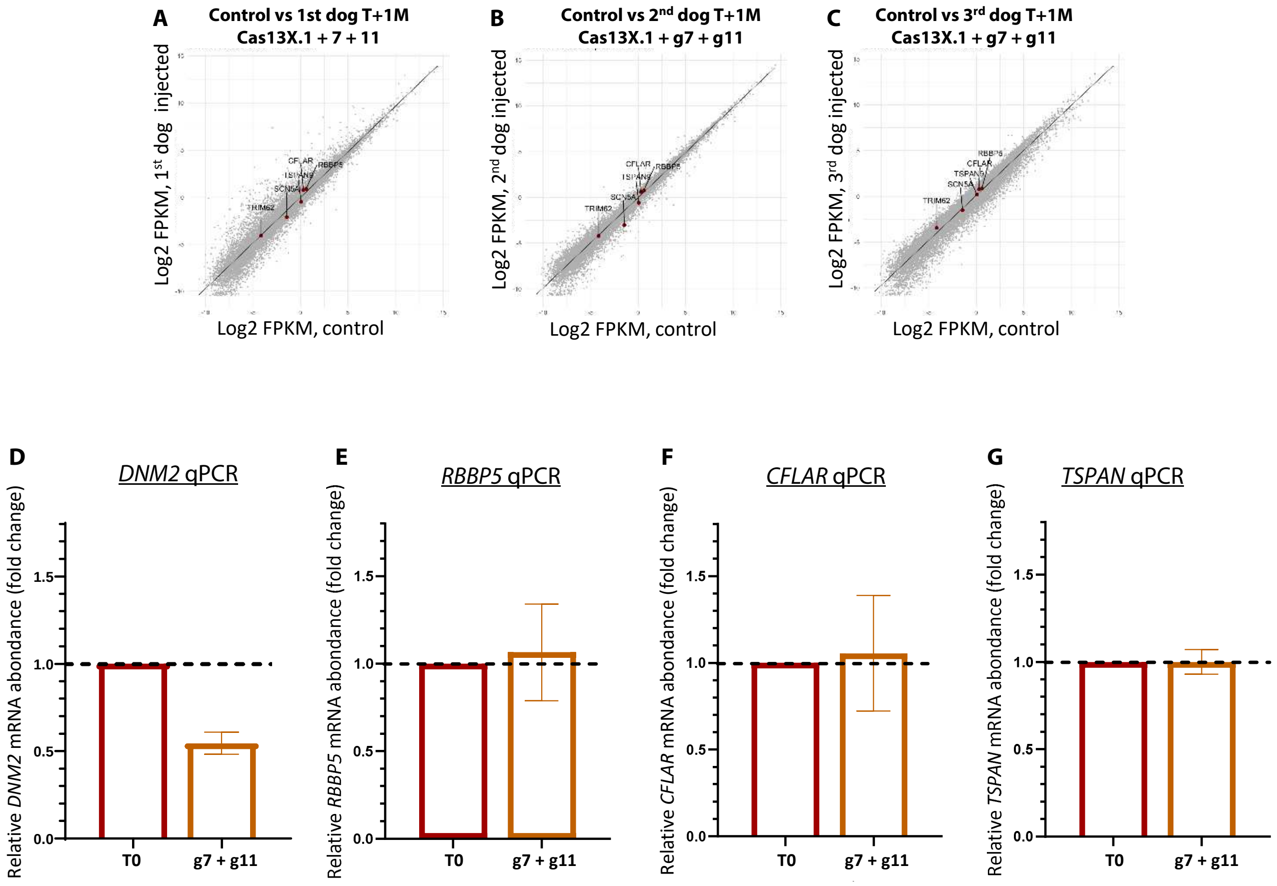


**Fig. S6: Absence of off-target effects in injected muscles: analysis by RNAseq and RT-qPCR.**

The results of the RNA-Seq analysis were presented in the form of scatter plots of the differential expression levels between the control biopsies (the three dogs aged 24 months before injection + biopsy at T0 of the first dog) and the biopsies taken after the treatment with the different conditions at $3.5x10^{12}$ vg - **A**: first dog injected with Cas13X.1 + g7 + g11 - **B**: second dog injected with Cas13X.1 + g7 + g11 - **C**: third dog injected with Cas13X.1 + g7 + g11. RT-qPCR analysis of (**D**) *DNM2*, (**E**) *RBBP5*, (**F**) *CFLAR* and (**G**) *TSPAN* transcripts. The mRNA was referred to *GAPDH* and *RPLS19* housekeeping genes and normalized results are expressed as mRNA relative fold change abundance related to the biopsies taken at T0.

**Table S1**

| **Name gRNA** | **Targeted sequence 5'-3'** | **Exon** |
|---|---|---|
| gRNA-1 | caaggacatgatcctgcagttcattagc | exon 4 |
| gRNA-2 | gaacaccttctccatggacccgcagcta | exon 19 |
| gRNA-3 | gtaccagatcaaggacatgatcctgcag | exon 4 |
| gRNA-4 | gatatcgagtaccagatcaaggacatga | exon 4 |
| gRNA-5 | gccgggagagcagcctcatcctcgccgt | exon 4 |
| gRNA-6 | gatggtccagcagtttggggtggacttt | exon 8 |
| gRNA-7 | aaggcttgcggaccatcggtgtcatca | exon 4/exon 5 |
| gRNA-8 | ggcgtggtgaaccgcagccagaaggaca | exon 6 |
| gRNA-9 | aacaccttctccatggacccgcagctag | exon 19 |
| gRNA-10 | aagaggctacattggcgtggtgaaccgc | exon 5/6 |
| gRNA-11 | ccaagctcgacctgatggatgaaggcac | exon 5 |

**Table S2**

| **Primer name** | **Sequence 5'- 3'** |
|---|---|
| DNM2- For | TACTGGTTTGTGCTGACAGC |
| DNM2- Rev | TGCTTGTTGGACATGAAGCC |
| RPLS19-For | CCTTCCTCAAAAAGTCTGGG |
| RPLS19-Rev | GTTCTCATCGTAGGGAGCAAG |
| GAPDH-For | TGGCAAAGTGGATATTGTCG |
| GAPDH-Rev | AGATGGACTTCCCGTTGATG |
| RBBP5 FOR | GGTCCAAACCTCCCTGATGG |
| RBBP5 REV | TGGATTTGCCATCCCCCTTC |
| CFLAR FOR | ACGACAGCTTTGTGTGTGTC |
| CFAR REV | ATGCGTCTCCCATGAACATC |
| TSPAN FOR | TCGTGTTTTACCTGGACCTGTC |
| TSPAN REV | ACAGCCACAGAGCCAGAATATC |
| LK0.1 | GACTATCATATGCTTACCGT |
| CasX.1 For | AAAGACCGCAGTGAACAAGG |
| CasX.1 Rev | TTCATCACGTCGCTGAACAG |
| crRNAg141 AS-RX | CTTG CCAAGCTCGACCTGATGGATGAAGGCAC |
| crRNAg8.16 AS-RX | CTTGAAGGCTTGCG GACCATCGGTGTCATCA |

**Supplementary Table 1 and Table 2 :**
List and sequences of the gRNAs (**1**) and the primers used (**2**)
